# Comparison of Genotypic and Phenotypic Correlations: Cheverud’s Conjecture in Humans

**DOI:** 10.1101/291062

**Authors:** Sebastian M. Sodini, Kathryn E. Kemper, Naomi R. Wray, Maciej Trzaskowski

## Abstract

Accurate estimation of genetic correlation requires large sample sizes and access to genetically informative data, which are not always available. Accordingly, phenotypic correlations are often assumed to reflect genotypic correlations in evolutionary biology. Cheverud’s conjecture asserts that the use of phenotypic correlations as proxies for genetic correlations is appropriate. Empirical evidence of the conjecture has been found across plant and animal species, with results suggesting that there is indeed a robust relationship between the two. Here, we investigate the conjecture in human populations, an analysis made possible by recent developments in availability of human genomic data and computing resources. A sample of 108,035 British European individuals from the UK Biobank was split equally into discovery and replication datasets. 17 traits were selected based on sample size, distribution and heritability. Genetic correlations were calculated using linkage disequilibrium score regression applied to the genome-wide association summary statistics of pairs of traits, and compared within and across datasets. Strong and significant correlations were found for the between-dataset comparison, suggesting that the genetic correlations from one independent sample were able to predict the phenotypic correlations from another independent sample within the same population. Designating the selected traits as morphological or non-morphological indicated little difference in correlation. The results of this study support the existence of a relationship between genetic and phenotypic correlations in humans. This finding is of specific interest in anthropological studies, which use measured phenotypic correlations to make inferences about the genetics of ancient human populations.

## Introduction

Genetic correlations are a measure of genetic factors shared between two traits. When two traits are highly genetically correlated, the genes that contribute to the traits are usually co-inherited (Lynch & Walsh, 1998). While traditionally used in animal breeding (Lynch & Walsh, 1998), in a broader research context, genetic correlations contribute to understanding the development and pathways of traits, population level gene flow and the co-occurrences of traits (Via & Hawthorne, 2005). For this reason, genetic correlations play an important role in evolutionary biology, and estimates of genetic correlations are also used in theoretical modelling of human populations.

Genetic correlations (*r*_*g*_) are calculated from the additive genetic variance and covariance between traits, as shown for traits X and Y, 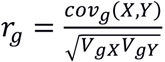, or for standardized traits where the phenotypic variances are one, 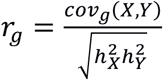, where 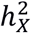and where 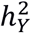are the heritability estimates of the two traits and *V*_*gX*_and *V*_*gY*_are the variances of the traits.

Traditionally genetic correlations are calculated from pedigree data using statistical methods to partition phenotypic (co)variance into genetic variance and genetic covariance (Henderson 1986). More recent methods make use of genome-wide single nucleotide polymorphism (SNP) data and the very small coefficients of relationship between very large numbers of unrelated individuals in order to calculate these parameters (Lee *et al.*, 2012). Since only common variants are included in the calculations, this approach assumes that the genetic correlation is the same across the allelic frequency spectrum. Accepting this caveat as reasonable, the approach has an advantage over the traditional methods, as unrelated individuals are less likely to have had exposure to similar environmental effects, reducing confounding from shared environment. Additionally, as genotyping becomes cheaper, genome-wide SNP data is becoming more readily and widely available than pedigree data. Moreover, unbiased estimates of genetic correlations are achievable with minimal computing resources from analysis of summary statistics from genome-wide association studies via the LD-score regression method (Bulik-Sullivan *et al.,* 2015a; Ni *et al.*, 2017).

The sampling variance of a genetic correlation estimate, depends on, and is larger than, the sampling variances of the concurrently estimated heritabilities (Robertson, 1959; Visscher *et al.*, 2014). Hence, large sample sizes are needed to estimate genetic correlations with accuracy. James Cheverud proposed in 1988 that phenotypic correlations (*r*_*p*_) could be used as a proxy for genetic correlations (Cheverud, 1988).

Whilst there has been criticism of the conjecture, most notably by Willis *et al.* (1991), subsequent studies in various organisms have provided much empirical evidence and theory supporting the conclusion. Roff (1996) considered a variety of traits from previously published datasets. This investigation showed that the relationship between the two correlations was most concordant in morphological traits, as opposed to behavioral or life history traits. In addition, while the average absolute disparity (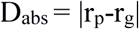 (Willis *et al.*, 1991)) between the correlations was relatively high (0.24- 0.46), this difference could be attributed to the sampling error of r_g_. In 2008, Kruuk *et al.* repeated the analysis of Roff’s 1996 paper with more recent data with an increased sample size, reaching similar conclusions.

The suitability of using phenotypic correlations as a proxy for genetic ones in various traits has been discussed by Hadfield *et al.* (2007), concluding that while the conjecture may be true in traits with high heritability, particularly those related to growth, there are still exceptions, and the conjecture most likely does not apply to all traits generally. Since phenotypic correlations depend both on the correlation of additive genetic and on the correlation of environmental effects (with the term environmental representing any effects that are not additive genetic), differences between phenotypic and genetic correlations must be explained by the relationship between genetic and environmental effects. Cheverud (1984) suggests that most environmental effects often act in the same direction and through the same pathways as genetic effects, which leads to a similarity between phenotypic and genetic correlations. Hadfield (2007), on the other hand, suggested that certain traits have environmental effects that act in the opposite direction to the genetic effects, which could reflect the conclusion of Roff (1996), who found lower correlation for life history and behavioral traits than morphological ones.

Despite the possible deficiencies of the conjecture when applied to non-morphological traits, behavioral researchers often assume that correlations between behaviors can give insight into the genetics behind the behavior. In order to test this assumption, Dochtermann (2011) tested the relationship between published behavioral genetic and phenotypic correlations from animal studies. The author found that while the correlation between the phenotypic and genetic correlations was high (r=0.86), the mean absolute difference between traits was also high (0.27), suggesting that phenotypic correlations were not a good predictor of genetic correlations between behavioral traits. Dochtermann found that while not a good predictor, the phenotypic correlation is able to reliably provide information on the direction of the genetic effect for behavioral research, still allowing to make certain genetic conclusions based on phenotypic data.

To date, studies investigating the existence of Cheverud’s conjecture in specific populations have looked at morphological traits in insects (Roff, 1995; Reusch & Blanckenhorn, 1998), tamarins (Ackermann & Cheverud, 2002), and plants (Waitt & Levin, 1998) with results corroborating the findings of Cheverud and Roff. While the conjecture has not been investigated in humans, it has been applied in human modelling. As genetic data are not directly accessible in many ancient human populations, phenotypic traits have been used to make conclusions regarding genetic information (Weaver *et al.*, 2007; Relethford & Blangero, 1990).

While the proportionality of phenotypic and genetic correlations has been assumed to be true in human populations, there has yet to be a study to investigate the conjecture in the context of humans. This study aims to fill the gap in understanding how the conjecture applies in human populations. Moreover, it aims to show whether human data differs from the results seen in animal and plant studies. Here, first we investigate the relationship between phenotypic and genetic correlations across 17 traits.

We then investigate the relationship by considering two general types of traits: morphological traits and other (non-morphological) traits. It was hypothesized that, similar to other species, genetic and phenotypic correlations are concordant in human traits, with a strong relationship particularly in morphological traits. Historically, the study of genetic correlations in humans has been limited by availability of data. This study uses genetic and phenotypic data, drawn from the first phase of the UK Biobank - a large sample of unrelated individuals (Sudlow *et al.*, 2015). This wealth of data allows a new look at Cheverud’s conjecture in the context of humans.

## Methods

Participants from the UK Biobank with British/Irish ancestry were selected based on self-reported ancestry and leading principal components calculated from SNP data, resulting in a sample size of 108,035 participants with available genotypes cleaned and imputed to a combined reference panel of 1000 Genomes and UK10K (see UKB documentation for details about QC and imputation, with sample selection following Robinson *et al.* (2017)). For our analyses we selected Hapmap3 SNPs, with minor allele frequency (MAF) > 0.01, a Hardy-Weinberg Equilibrium (HWE) test p-value > 1.0E-6 and imputation info-score > 0.3. The total sample was randomly split into two sets (n=54,017, n=54018), with no evidence for differences in demographic variables (Supplementary Table 1). This allowed us to estimate genetic and phenotypic correlations within in each set, and also allowed estimation of genetic correlations between the two independent sets.

**Table 1:** *Final list of traits used in study with corresponding sample size for both discovery and replication samples.*

| **Morphological Traits** |  | **Non-Morphological Traits** |  |
| --- | --- | --- | --- |
| **Trait** | **Sample Size\*** | **Trait** | **Sample Size\*** |
| Body Mass Index | 53871/53867 | Basal Metabolic Rate | 53112/53087 |
| Body Fat Percentage | 53086/53046 | Diastolic Blood Pressure | 50801/50682 |
| Forced Vital Capacity | 42341/42336 | Heel Bone Density | 31254/31174 |
| Height | 53931/53926 | Neuroticism Score | 43940/44204 |
| Hip Circumference | 53940/53937 | Pulse Rate | 50801/50682 |
| Peak Expiratory Flow | 48399/48262 | Reaction Time | 53693/53716 |
| Waist Circumference | 53942/53941 | Systolic Blood Pressure | 50801/50682 |
| Weight | 53886/53885 |  |  |
| Grip Strength (R) | 53802/53789 |  |  |
| Grip Strength (L) | 53803/53796 |  |  |
| **Average** | **48927/48897** | **Average** | **48927/48897** |

\* Formatted as “discovery/replication”

Traits with > 10,000 observations in each dataset were selected for analysis. Selection of these traits included inspecting the distribution, and traits with drastically non-normal distributions were excluded. Key covariates and exclusion variables were identified for all traits. Exclusions were handled on a trait-by-trait basis. For example, subjects were excluded from analysis for spirometry traits if they had smoked within the last hour (see Supplementary Table 2). The effects of sex, age, age^2^ and testing centre were regressed out of the data using a linear model. Traits relating to the cardiovascular system had the effect of blood pressure medication regressed out (medication use was taken as a binary variable). Genetically derived principal components were also used as covariates, however only when calculating genetic correlations, and not phenotypic ones. This was done to emulate a situation where genetic information is not available, which is where Cheverud’s conjecture is relevant. Finally, the residuals were transformed with a rank normal transformation (Van der Waerden transformation; (Lehaman, 1975)).

**Table 2:** Quantitative comparisons of phenotypic and genetic correlations

|  | Discovery genetic<br>Replication phenotypic |  |  |  | Discovery phenotypic<br>Replication genetic |  |  |  |
| --- | --- | --- | --- | --- | --- | --- | --- | --- |
|  | r | Slope | Intercept | Avg D<sub>abs</sub>‡ | r | Slope | Intercept | Avg D<sub>abs</sub>‡ |
| Morphological | 0.97*(0.03) | 1.08† | 0.01 | 0.09(0.04) | 0.97*(0.03) | 1.03 | 0.04 | 0.08(0.04) |
| Non-Morphological | 0.93*(0.06) | 0.92 | -0.01 | 0.06(0.07) | 0.93*(0.06) | 0.90 | 0.00 | 0.05(0.06) |
| Morphological/Non-morphological | 0.96*(0.03) | 1.08 | -0.01 | 0.05(0.05) | 0.96*(0.03) | 1.10 | -0.03 | 0.05(0.05) |
| Combined | 0.97*(0.02) | 1.09† | -0.01 | 0.06(0.05) | 0.97*(0.02) | 1.07† | 0.00 | 0.06(0.05) |

|  | Discovery |  |  |  | Replication |  |  |  |
| --- | --- | --- | --- | --- | --- | --- | --- | --- |
|  | r | Slope | Intercept | Avg D<sub>abs</sub>‡ | r | Slope | Intercept | Avg D<sub>abs</sub>‡ |
| Morphological | 0.97*(0.03) | 1.09† | 0.01 | 0.09(0.04) | 0.97*(0.03) | 1.03 | 0.04 | 0.08(0.04) |
| Non-Morphological | 0.93*(0.06) | 0.92 | -0.01 | 0.05(0.07) | 0.92*(0.06) | 0.89 | 0.00 | 0.05(0.06) |
| Morphological/Non-morphological | 0.96*(0.03) | 1.07 | -0.01 | 0.05(0.05) | 0.96*(0.04) | 1.10 | -0.03 | 0.06(0.05) |
| Combined | 0.97*(0.02) | 1.09† | -0.01 | 0.06(0.05) | 0.96*(0.02) | 1.07† | 0.00 | 0.06(0.05) |

\*=significant at $p < 0.003125$ (Bonferroni multiple testing correction)
†=significant difference from unity line ( $p < 0.003125$ )
‡=average of absolute disparity

Phenotypic correlations were estimated as Pearson correlations between each pair of traits, within both discovery and replication data sets (Figure 1). A genome-wide association study (GWAS) analysis was performed using PLINK 1.9 (Chang *et al.*, 2015) for each trait in discovery and replication samples separately, using a linear association model. The proportion of variance attributable to genome-wide SNPs (SNP-heritability) and the genetic correlation attributable to genome-wide SNPs was estimated from the GWAS summary statistics using an LD-score regression analysis as implemented by Bulik-Sullivan *et al.* (2015b) in the LDSC software package, using LD-scores estimated from the full data set. Briefly, genetic variances (or covariances) are estimated as a function of regressions of the square (or product) of association analysis z-statistics of SNPs for traits (or pairs of traits) on their linkage disequilibrium scores (LD-scores), where an LD-score is the sum of LD r^2^ made by the SNP with all other SNPs. The method assumes that traits have a polygenic genetic architecture. LD-score estimates of genetic correlations agree well with those based on mixed model analysis of full individual level genotype data (e.g., GREML in GCTA; Yang *et al.*, 2010; Lee *et al.,* 2012), but are achieved at a small fraction of computing resources, albeit with higher standard errors (Bulik-Sullivan *et al.,* 2015a; Ni *et al.*, 2017). Traits with estimated SNP-heritability less than 0.05 were removed, as the estimates of genetic correlation are unstable for traits with low SNP-heritability. Seventeen traits were used in the final analysis (Table 1), which were characterized as either morphological (n=10) or non-morphological (n=7), thereby generating 45 pairwise correlations within the morphological traits, 21 between non-morphological traits and 70 correlations between-traits for each data set. Genetic correlations were also estimated between all pairs of traits between the two data sets.

**Figure 1:**
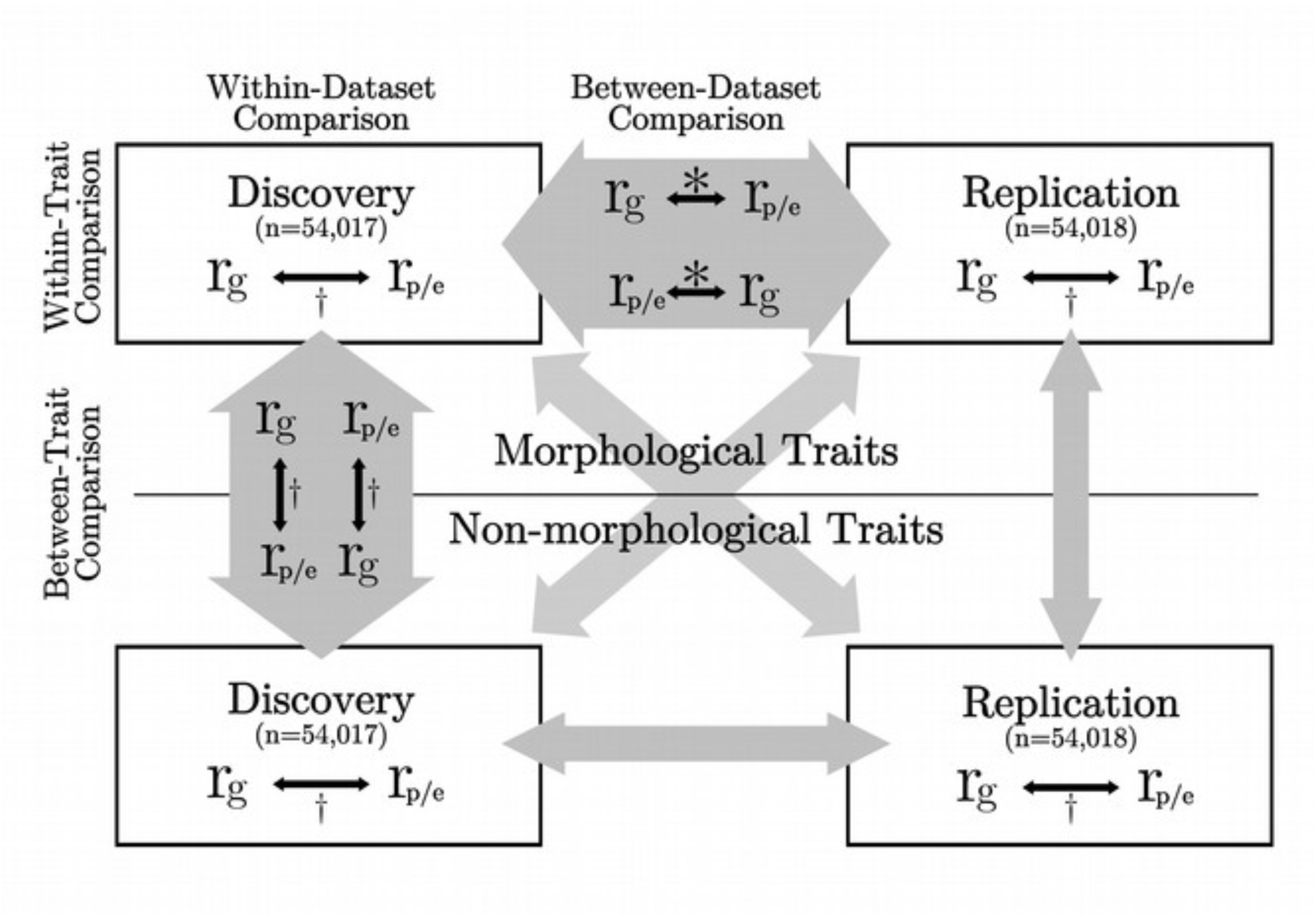
Schematic diagram of statistical analyses performed. 108,035 British European individuals were evenly divided into discovery and replication datasets. Genetic and phenotypic correlations were calculated within group for 17 traits. Black arrows show the comparisons performed. Empty grey arrows indicate comparisons similar to the equivalent grey arrow (ie. the within-replication, between-trait comparison is the same as the within-discovery, between-trait comparison). *=Figure 3, Table 2A †= Table 2B

Pearson correlation coefficient, linear regression and absolute disparity (Willis *et al.*, 1991), were calculated for within-trait, between-trait and all traits (combined) in both within-dataset and between-dataset comparisons (see Figure 1). The difference of the slope from the unity line was assessed by comparing the least squares linear regression to a linear model with a slope of one. Significance of the slope being different from one was set at p < 0.003125, with Bonferroni correction for 16 tests.

Comparisons of the environmental correlations (r_e_) and genetic correlations were also performed,where 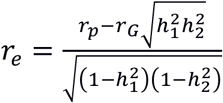(Supplementary Text 1). Similar analyses were performed as with the phenotypic correlation, but using the environmental correlation in its place. The results of the analysis are shown in Supplementary Figure 1 and Supplementary Table 3.

Finally, in sensitivity analyses to assess the similarity of the structure of the matrices, various matrix similarity tests were applied, as discussed by Roff *et al.* (2012). It is suggested that a variety of these tests should be used, as it is possible that they are not all sensitive to the same differences between matrices. The random skewers, T-test and T^2^-test, and modified Mantel test were applied to compare phenotypic and genetic correlations. The random skewers method investigates whether two matrices respond similarly to selection (Cheverud & Marroig, 2007), the T-test and T^2^-test consider the equality by examining the sum of the absolute difference or squared difference between matrix elements, and the modified Mantel test looks at the correlation between the matrix elements. Results for each of the tests are shown in Supplementary Materials - Table 4.

Given the sample sizes available, phenotypic correlations were estimated with high accuracy. There is no current literature on the expected standard error or power from LD score regression, however it can be compared to those expected from the linear mixed model maximum likelihood method (GREML), which estimates SNP-heritabilities and genetic correlations from GWAS genotype data (Visscher *et al*., 2014). Empirical comparisons have shown that the error associated with using LD score regression is approximately fifty percent larger than that of GREML (Ni *et al.*, 2017). Using the GCTA-GREML power calculator developed by Visscher *et al*., the trait with the smallest sample size (heel bone density, n=31254/31174) has a power of “0.99” to detect the heritability cutoff of 0.05, with a standard error of 0.0101. The pair of traits with lowest sample size (heel bone density and forced vital capacity) had a power of 0.98, and a standard error of 0.0219 to detect the genetic correlation of −0.089, as estimated by LDSC. In comparison, the observed standard error from LDSC was 0.051, a little more than double that predicted for bivariate GREML, although still relatively low. Hence, we conclude that the UK Biobank Pilot data is well-powered for the analyses conducted.

## Results

Across all traits the estimated SNP-heritabilities ranged from 0.073 to 0.52, with a mean of 0.20 (Figure 2). The standard errors of the heritability estimates reflected the sample sizes and ranged from 0.009 to 0.042. Morphological traits had a higher average estimated SNP-heritability (0.23) than non-morphological traits (0.16), but the difference was not significant (p=0.22). The Pearson correlation coefficients between the phenotypic and genetic correlations for the combined comparison of all 17 traits were r=0.97 and r=0.96 for each of the between-dataset comparisons (Table 2A). The least squares linear regression coefficient was significantly different from the unity line when considering all traits combined, however it was not significant when considering only morphological or non-morphological traits (Table 2A). The mean difference between correlations was 0.06 in both cases, calculated using the method described by Willis *et al.* (1991) and described earlier (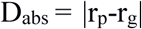). This difference was not significantly different from 0 for both discovery and replication datasets. The maximum difference between two correlations was 0.24 and the minimum was 0.0004. On average, the magnitude of genetic correlations was 0.04 higher than phenotypic ones.

**Figure 2:**
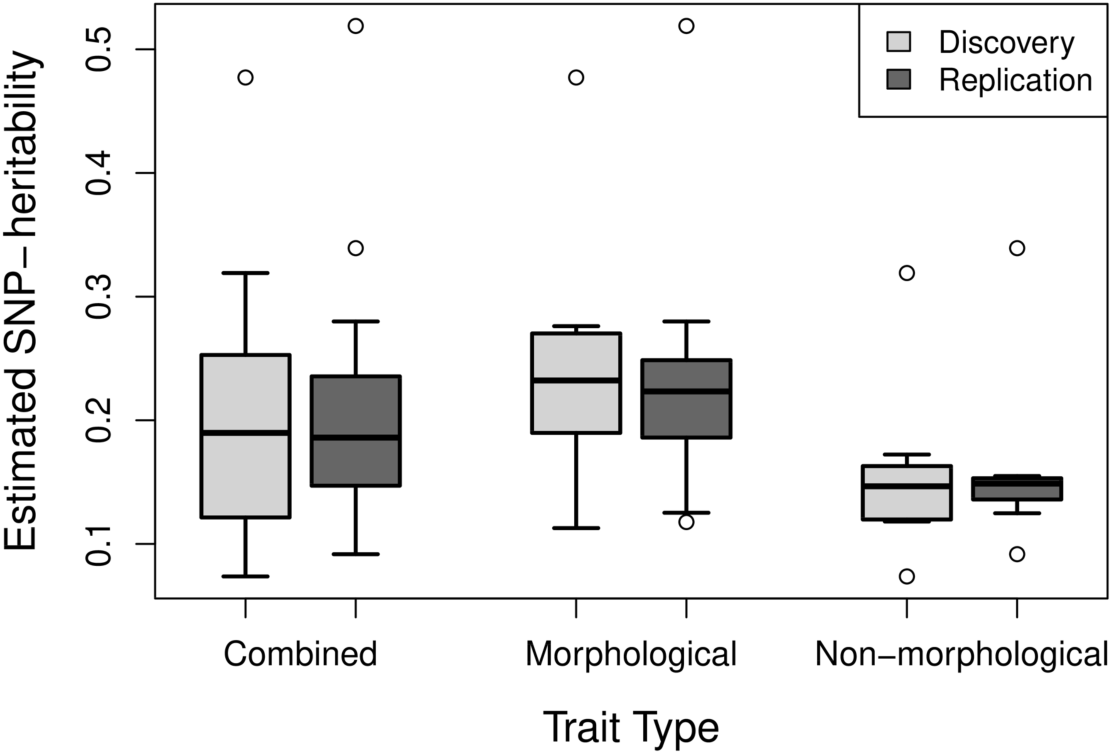
Boxplots of the distribution of estimated SNP-heritabilities for all traits (combined, 17 traits), morphological traits (10 traits), and non-morphological traits (7 traits). Quantitative traits were selected from the UK Biobank and SNP-heritabilities estimated through LD-score regression. Sample sizes used to calculate SNP-heritabilities range from 31174 to 53942 individuals.

Comparison between morphological and non-morphological traits showed some general differences between the two types of traits. Both types of traits had strong positive correlations across both datasets (between r=0.92 and r=0.97, Table 2A). However, the distribution of correlations was different between the two groups: morphological traits were normally distributed with a range of genetic and phenotypic correlations (between 0 and 1) while distribution of the non-morphological trait correlations was right-hand skewed with a mean closer to 0 (Figure 3). In both sets of traits, however, least squares linear regression was not significantly different from the unity line. The mean absolute disparity between correlations ranged between 0.05 and 0.09 (Table 2A). Very similar results to those above were seen in within-dataset analysis (Table 2B). None of the parameters changed appreciably, and any differences were lost when rounding values.

**Figure 3:**
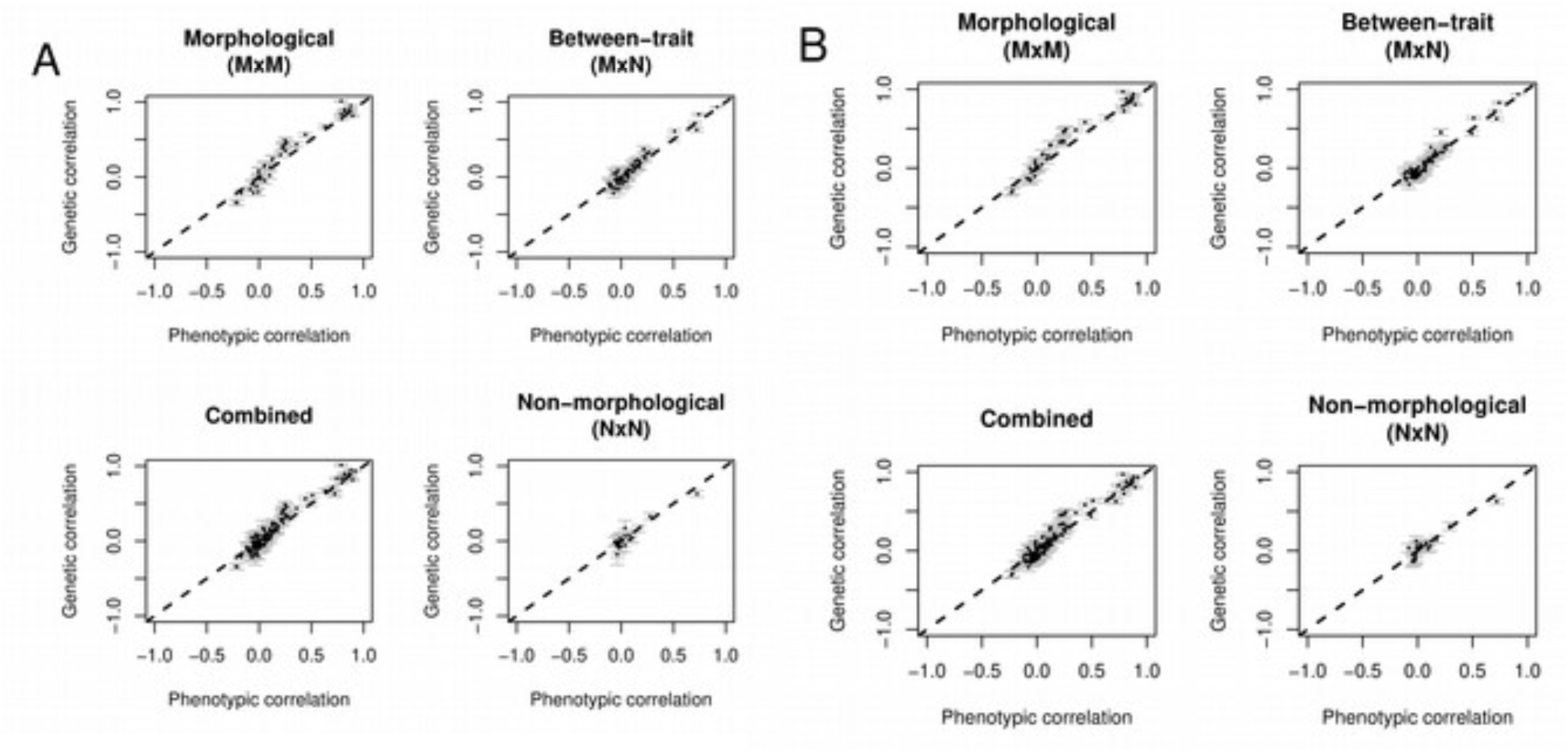
Plots of genetic correlation versus phenotypic correlation for the between-dataset comparison. 108,035 British European individuals were distributed into discovery (n=54017) and replication (n=54018) datasets. Genetic and phenotypic correlations were calculated within group for 17 traits. **A.** Genetic correlations from discovery dataset, phenotypic correlations from replication dataset. **B.** Genetic correlations from replication dataset, phenotypic correlations from discovery dataset. The between-trait comparison refers to the correlations between morphological (M) and non-morphological traits (N).

Repeating the same analysis with environmental correlation showed a similar result, albeit with slightly lower levels of correlation (r=0.90-0.96), and slightly higher mean absolute disparity (0.06- 0.11). Of note, non-morphological comparisons were lower than the morphological comparison for the within-trait correlation in the discovery dataset. Full results can be found in Supplementary Table 3.

Comparison between the phenotypic and genetic correlations using the random skewers method had p-values of 1.0 for all comparisons, giving no evidence to reject the null hypothesis (Supplementary Material – Table 4). Both the T-test and T^2^-test comparisons showed no overall difference between the off-diagonal elements of the matrices, and the modified Mantel test had a p-value of 1.0, supporting the null hypothesis of correlation between the matrix elements (Roff *et al.*, 2012). Plots of difference in correlation versus mean heritability and mean sample size, as well as standard error versus mean heritability and mean sample size can be found in Supplementary Figure 2.

## Discussion

The aim of this study was to investigate the relationship between genetic and phenotypic correlations in humans using data from large samples of unrelated individuals (ie very distantly related) from the UK Biobank. Based on reports from other species, we hypothesized a strong correlation between genetic and phenotypic correlations but with a stronger correlation between morphological than non-morphological traits. Our analyses confirmed these hypotheses, but it is notable that the phenotypic and genetic correlations between non-morphological traits, while often different from zero, were smaller than those between morphological traits. High Pearson correlation coefficients were seen across both of the between-dataset comparisons (0.92-0.97), as well as in within-dataset correlations (0.93-0.97) (Table 2A,B). These findings indicate that the results are reproducible in independent samples, and more practically, that overall the phenotypic correlations from one group are good predictors of genetic correlations in an independent sample of the same ethnicity. The mean absolute disparity between the combined phenotypic and genetic correlations was not significantly different from zero in both between-dataset comparisons (Table 2), as well as in within-dataset comparisons (Table 3). These values support the conclusion of Roff (1996) and Kruuk *et al*. (2008) who suggested that their reported differences (0.24-0.46 and 0.245 respectively) are a reflection of sampling error of r_g_. The mean absolute disparity in the current paper is much lower, reflecting the larger sample sizes lowering the sampling error of r_g_. Additionally, application of the random skewers, T-test, T^2^-test and modified Mantel test methods (Roff *et al.*, 2012; Cheverud & Marroig, 2007) indicated similar structure between the covariance matrices (Supplementary Material 4). In conclusion, these results confirm the prior assumptions used in anthropometric studies. Just as it was true in other species, phenotypic correlations are good proxies for genetic correlations in human traits.

Comparison between morphological and non-morphological traits showed little difference between the two in terms of the relationship between phenotypic and genetic correlations, although there was a difference in the average magnitude of the correlations (mean magnitude of genetic correlations between morphological traits was 0.39(SE=0.04) and between non-morphological traits was 0.11(SE=0.07), difference p=5×10^−5^). While the correlation coefficient of the morphological within-trait, between-dataset comparison (r=0.97/0.97, Table 2A) was higher than that of the non-morphological comparison (r=0.93/0.92, Table 2A), this was not a significant difference. This finding was also true in the within-dataset comparisons. It is possible that this difference in correlation is driven by the difference in SNP-heritability (Figure 2), and thus accuracy of r_g_ estimation. However, while geometric mean heritability of the pair of traits and the standard error of r_g_ is negatively correlated (Supplementary Figure 2), this is not true between the mean heritability and the difference between the phenotypic and genetic correlations (Supplementary Figure 2). This would suggest that the difference in SNP-heritability between the traits does not play a major role in the differences between phenotypic and genetic correlations, and thus does not contribute to the differences between morphological and non-morphological traits seen in this study. The between-trait, between-dataset comparison showed high correlation between the two types of traits (Table 2, Figure 3). It is worth noting that the strong overall phenotypic correlation between morphological and non-morphological traits may be a characteristic of the non-morphological traits selected in this study. The selected traits may not be representative of the whole spectrum of non-morphological traits. Consequently, the relationship of other non-morphological traits could be different from that observed here.

To summarise, a strong correlation of phenotypic and genetic correlations was found in human traits. This finding is novel in the context of humans, as previous analyses of this kind were limited by sample size and techniques. Additionally, the correlation relationship between phenotypic and genetic correlations was consistent between morphological and non-morphological traits. This is a surprising result given previous literature in the area, which suggested that morphological traits may fit the conjecture better than life history traits (Roff, 1996), but as discussed, this could partly be due to the traits selected for this study as the non-morphological traits are not representative of life history traits. Additionally, the distinction between the categories of “morphological” and “non-morphological” is unclear for some traits. For example, forced vital capacity is directly related to lung volume, a morphological trait. On the other hand, non-morphological factors such as lung compliance, muscle strength and mucus secretions also affect the forced vital capacity, making it difficult to classify the trait. While the phenotypic and genetic correlations between non-morphological traits were relatively low, those between morphological and non-morphological traits covered a similar range, but had on average a slightly lower magnitude than those between the morphological traits. An important assumption of our approach is that the genetic correlations estimated from genome-wide SNP data are representative of the genetic correlations of variants across the allelic spectrum, but this seems to be a reasonable assumption. For example, the genetic correlation estimate calculated in this paper between BMI and body fat percentage was 0.86, consistent with estimates from twin studies (Faith *et al.,* 1999). Another example is the correlation between systolic blood pressure and BMI, which was 0.21 in this study, consistent with twin studies (Cui *et al.*, 2002).

The biological mechanism for the expected difference between types of traits discussed by Waitt & Levin (1998) is that of phenotypic plasticity - additive environmental effects on a trait. One of the criticisms of Cheverud’s conjecture by Willis *et al*. (1991) was that most of the data used in the original paper came from laboratory grown animals, leading to an underestimation of the environmental effects, which would be found in nature. Cheverud (1984) suggested that environmental and genetic effects are governed by the same developmental constraints and thus should have similar patterns, decreasing the impact on the correlation between traits. Hadfield *et al.* (2007), on the other hand, suggested that for certain groups of traits, the genetic and environmental effects act in opposing directions, decreasing correlation. In this study, r_g_ and r_e_ were positively correlated (r=0.90-0.96, Supplementary Table 3), suggesting that the genetic and environmental effects have similar correlational patterns. This provides support for Cheverud’s suggestion, and overcomes the “underestimation of environmental effects” argument posed by Willis et al. (1991), as the UK Biobank is a population based community sample. However, in the case of non-morphological traits, it may also indicate that our sample of traits is not fully representative of the whole spectrum.

When calculating the genetic correlations, many covariates were used in order to best estimate the value of r_g_, including genetically derived principal components. In order to simulate a scenario where no genetic information is available, phenotypic correlations did not include covariates that contained genetic information. Instead, they were limited to covariates that would have been available without such information (age, age^2^, sex and location). When these covariates were not accounted for, the mean absolute disparity ranged between 0.06 and 0.20 (Supplementary Table 5), higher than when covariates are accounted for (Table 2). Whilst the disparity without covariates is still quite low compared to prior literature, this finding indicates that environmental effects do play a role in modulating the phenotypic correlation, as suggested by phenotypic plasticity. Thus, it is important to account for some of the major confounding effects when using phenotypic correlations to estimate genetic ones, although in some studies confounding factors may not be recorded.

The results of this study are of specific interest in anthropological studies where anthropometric measurements are used as a proxy for genetic information. The results presented show support for this approximation in human studies, although, care should be taken when extrapolating the results of this study to other populations and environmental contexts, such as in ancient human populations subject to anthropological studies. The evidence provided here is based on observations in modern human populations, which may differ from earlier human populations. For example, large-scale famine and infections would have often affected earlier human populations, but are less of an issue for modern Europeans. Despite this, it is often already assumed that the phenotypic and genetic variance-covariance matrices are proportional between modern humans and even Neanderthals (Weaver *et al.*, 2007). Another caveat is that the morphological traits used in this study differ from those used in anthropometric studies. Nonetheless, the evidence from this study suggests that morphological traits do appear to fit Cheverud’s conjecture well, supporting its use in these kinds of traits.

In conclusion, this study investigated Cheverud’s conjecture in the context of human genetics. Correlations calculated using LD-score regression utilizing data from the UK Biobank support the validity of the conjecture in human populations. This study provides the quantitative evidence to support the use of phenotypic correlations as a proxy for genetic correlations in studies where genetic information is not available.

## Acknowledgements

We acknowledge funding from the National Health and Medical Research Council (grants (1078901, 1087889, 1113400)). We thank Katarina McGuigan for providing feedback on this work. This research has been conducted using the UK Biobank Resource under project 12514.

## Supplementary Material

**Supplementary Table 1.** **Population Demographic variables between samples**

| Trait | Discovery (mean(sd)) | Replication (mean(sd)) | Percent Difference (%) |
| --- | --- | --- | --- |
| Number Male | 28311 | 28452 | 0.50 |
| Number Female | 25706 | 25566 | -0.54 |
| Age | 56.91(7.1) | 56.93(7.9) | 0.04 |
| Height | 168.85(9.2) | 168.82(9.2) | -0.02 |
| Weight | 78.68(16.0) | 78.68(16.1) | -0.007 |
| BMI | 27.52(4.8) | 27.52(4.8) | 0.02 |

108,035 British European individuals were distributed into discovery and replication datasets. The above table shows the distribution of certain demographic characteristics between the two datasets. Overall, there is only a small difference in the demographics between the datasets.

**Supplementary Table 2.**
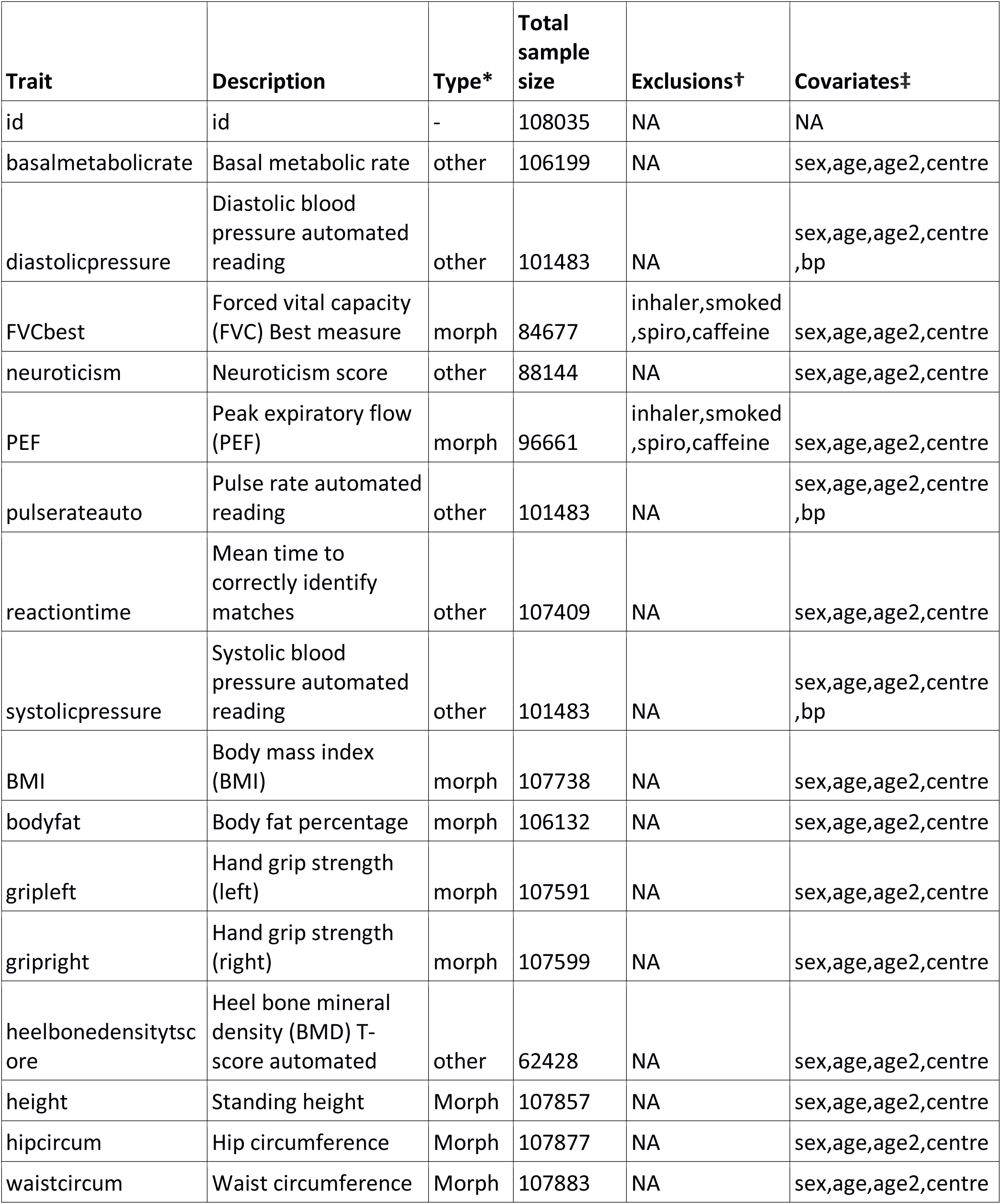
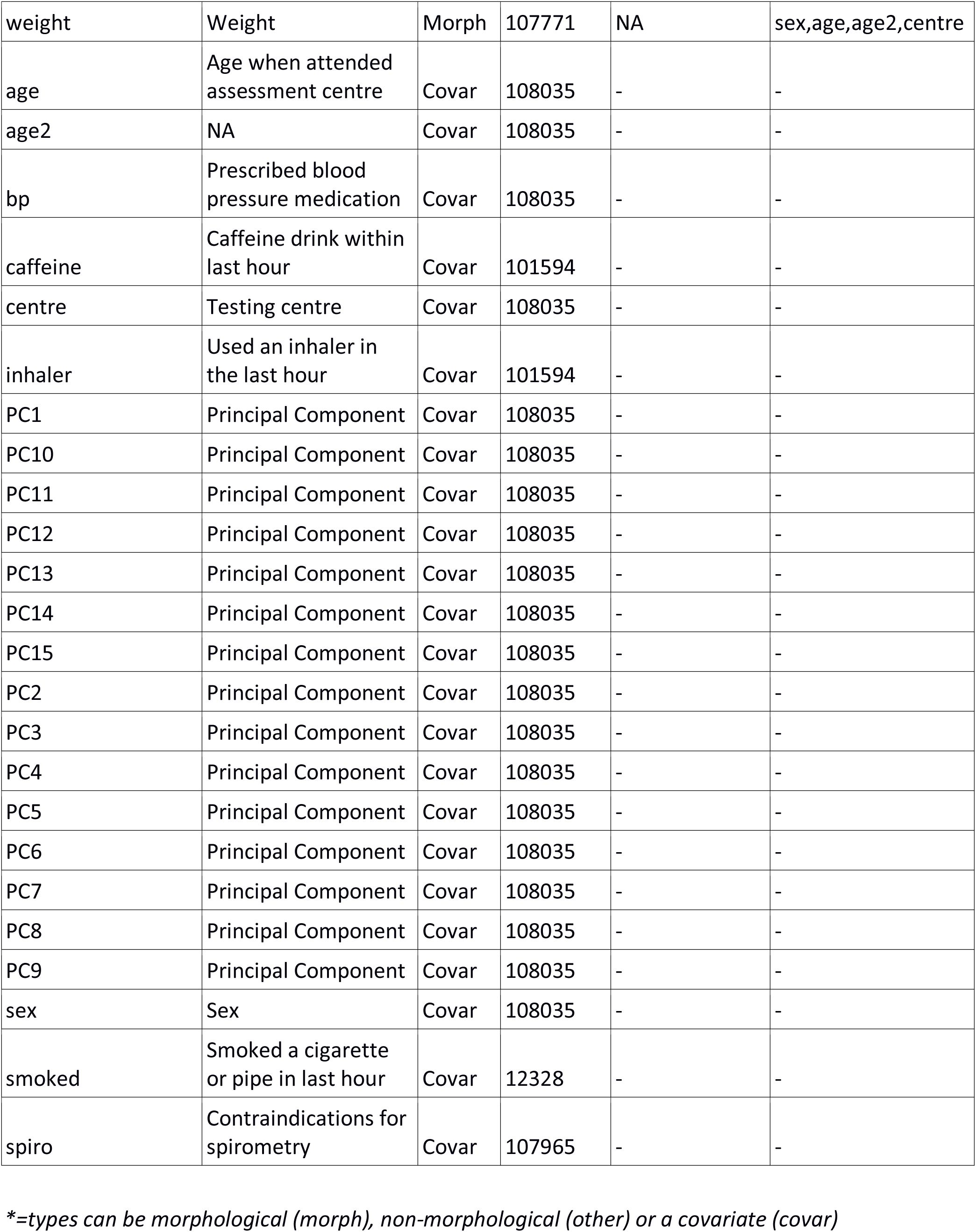
All traits with exclusions and covariates.

### Supplementary Text 1

Derivation of r_e_ Within each dataset there is 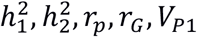 and *V*_*P*2_.
Since:

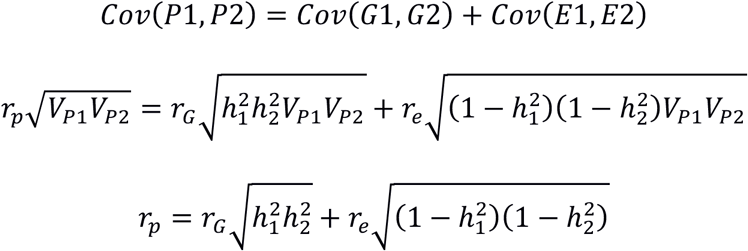

And thus:

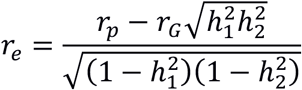

**Supplementary Figure 1.**
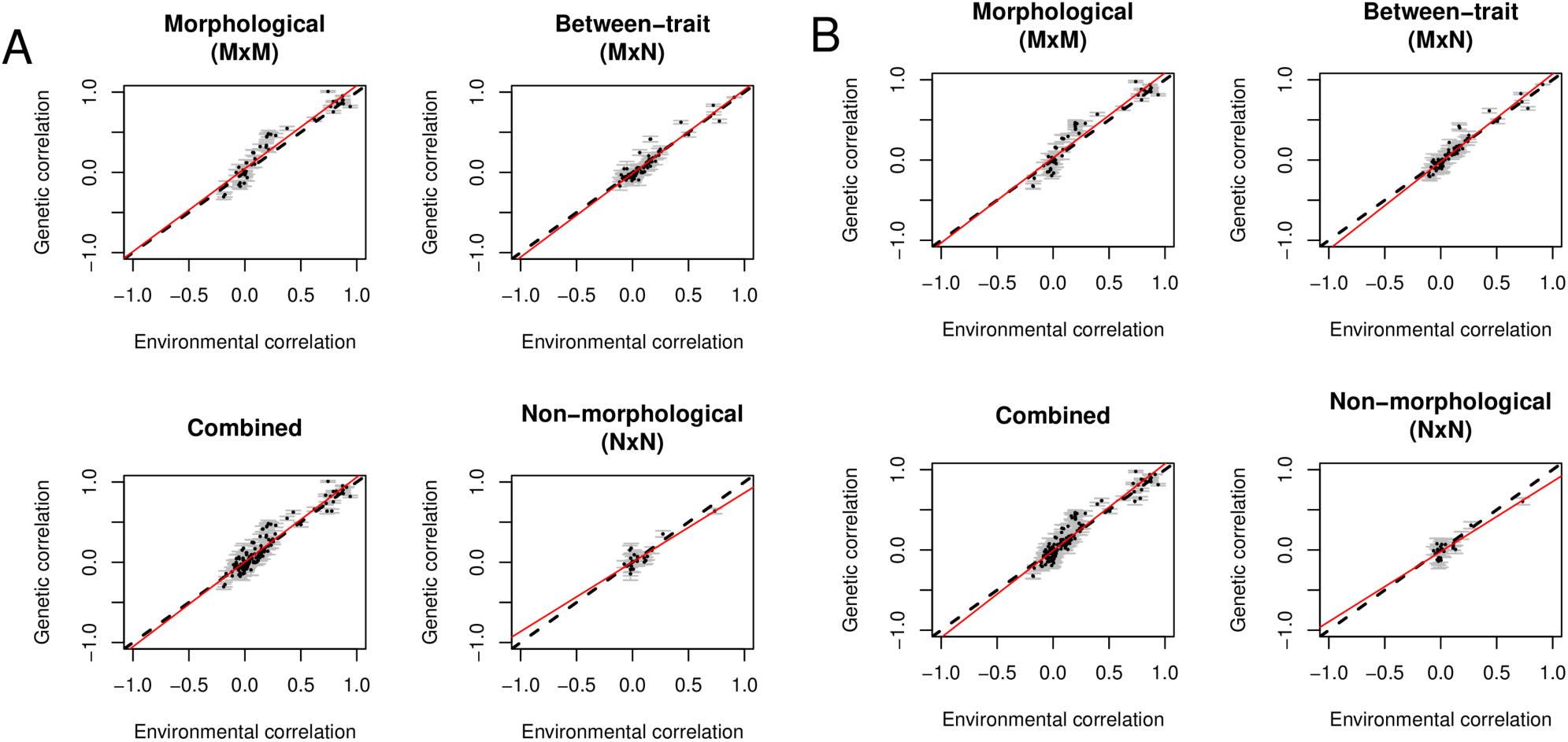
Comparison plots of between-dataset r_g_ and r_e_. Plots of genetic correlation versus environmental correlation for the between-dataset comparison. 108,035 British European individuals were evenly distributed into discovery and replication datasets. Genetic and phenotypic correlations were calculated within group for 17 traits. Environmental correlations were calculated using the formula 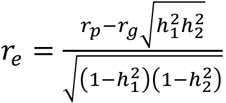, as derived in the supplementary text. **A.** Genetic correlations from discovery dataset, environmental correlations from replication dataset. **B.** Genetic correlations from replication dataset, environmental correlations from discovery dataset.

**Supplementary Table 3.** Summary statistics of linear regression between r_g_ and r_e_ for between-dataset and within-dataset comparisons. The following tables contain the summary statistics of least squares linear regression of environmental correlations and between-dataset genetic correlations for all traits (combined), morphological traits, and non-morphological traits. Table headings indicate which correlations are being compared between the groups. Environmental correlation was calculated using the formula 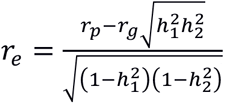, as derived in the supplementary text.

|  | Discovery genetic<br>Replication environmental |  |  |  | Discovery environmental<br>Replication genetic |  |  |  |
| --- | --- | --- | --- | --- | --- | --- | --- | --- |
|  | r | Slope | Intercept | Avg D<sub>abs</sub>‡ | r | Slope | Intercept | Avg D<sub>abs</sub>‡ |
| Morphological | 0.95*(0.03) | 1.08 | 0.02 | 0.11(0.04) | 0.96*(0.03) | 1.03 | 0.05 | 0.10(0.04) |
| Non-Morphological | 0.91*(0.07) | 0.89 | -0.01 | 0.06(0.07) | 0.90*(0.07) | 0.86 | 0.00 | 0.06(0.06) |
| Between-trait | 0.95*(0.04) | 1.06 | -0.01 | 0.06(0.05) | 0.94*(0.04) | 1.08 | -0.03 | 0.06(0.05) |
| Combined | 0.95*(0.02) | 1.08† | 0.00 | 0.08(0.05) | 0.95*(0.02) | 1.07† | 0.00 | 0.08(0.05) |

|  | Discovery |  |  |  | Replication |  |  |  |
| --- | --- | --- | --- | --- | --- | --- | --- | --- |
|  | r | Slope | Intercept | Avg D<sub>abs</sub>‡ | r | Slope | Intercept | Avg D<sub>abs</sub>‡ |
| Morphological | 0.95*(0.03) | 1.08 | 0.02 | 0.11(0.04) | 0.95*(0.03) | 1.02 | 0.05 | 0.10(0.04) |
| Non-Morphological | 0.89*(0.07) | 0.87 | -0.01 | 0.07(0.07) | 0.90*(0.07) | 0.86 | 0.00 | 0.06(0.06) |
| Between-trait | 0.94*(0.04) | 1.05 | -0.01 | 0.06(0.05) | 0.94*(0.04) | 1.08 | -0.03 | 0.06(0.05) |
| Combined | 0.95*(0.02) | 1.08† | 0.00 | 0.08(0.05) | 0.95*(0.02) | 1.06 | 0.00 | 0.08(0.05) |

\*=significant at $p < 0.003125$ (Bonferroni multiple testing correction)
†=significant difference from unity line ( $p < 0.003125$ )
‡=average of absolute disparity

**Supplementary Table 4.**
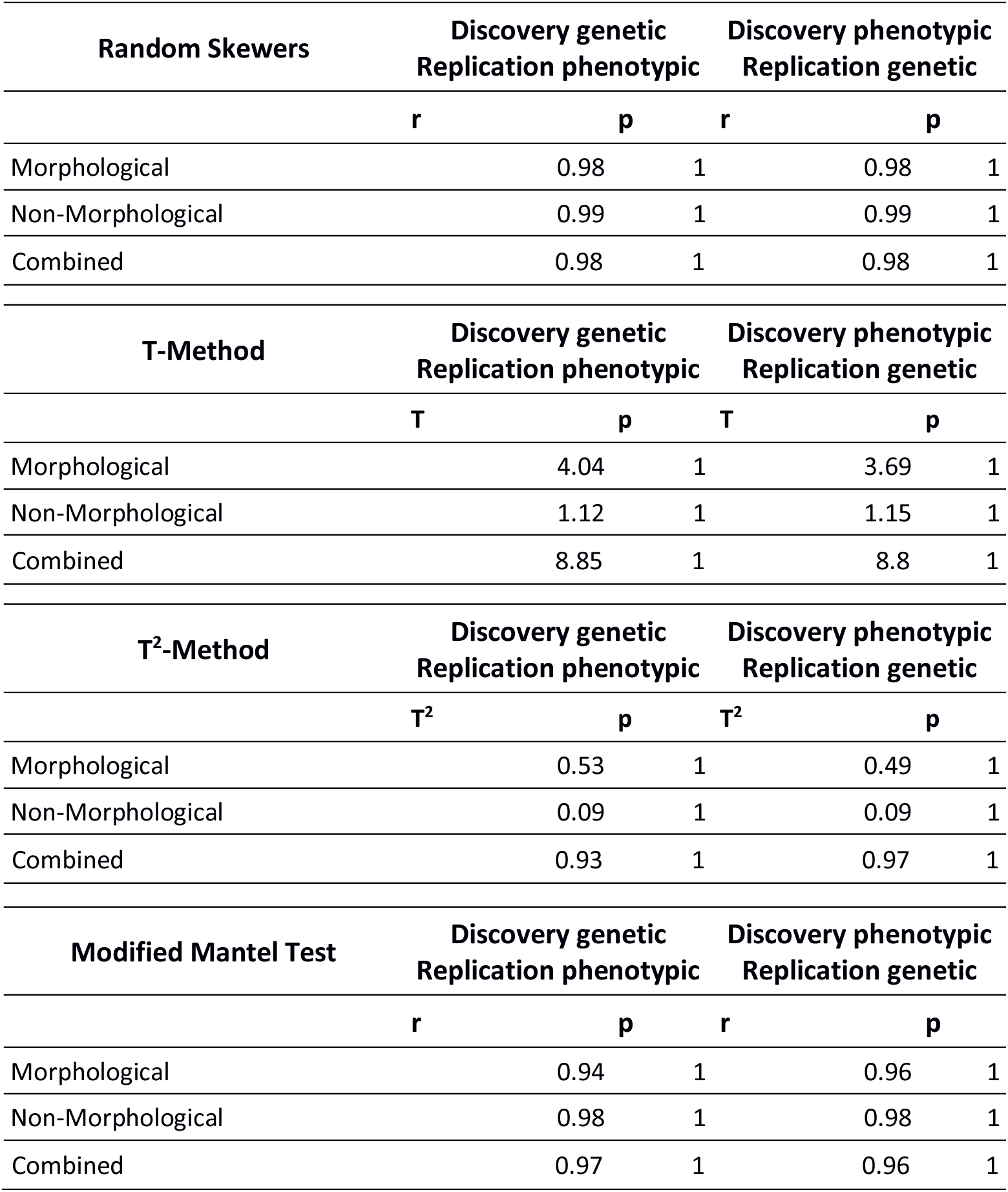
Summary of results from matrix comparison methods. The comparison methods were implemented as supplied by Roff, 2012 (doi: 10.5061/dryad.kb27f3t1), adapted to compare off-diagonal of the matrices. The null hypothesis for each test is similarity between the matrices.

**Supplementary Figure 2.**
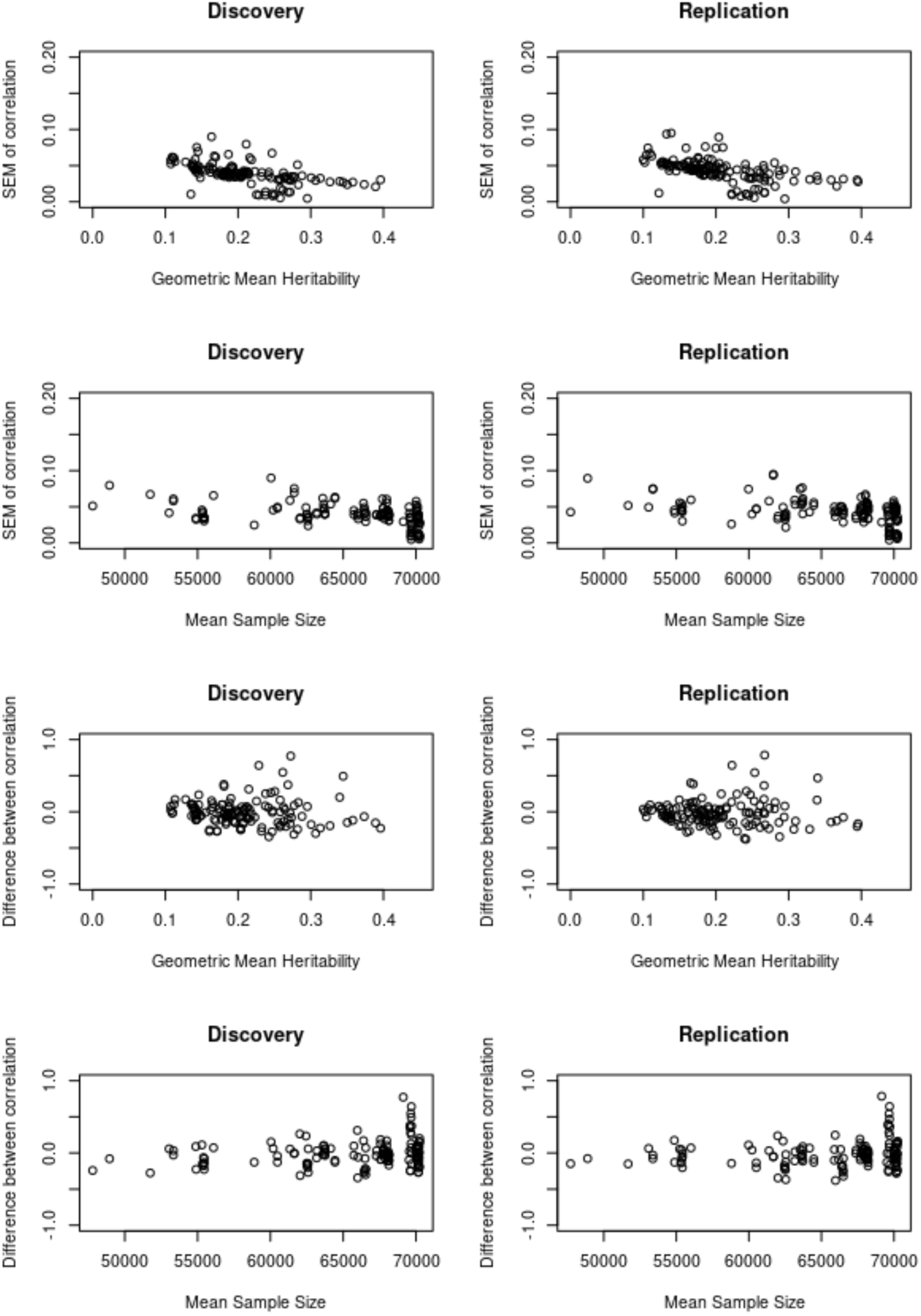
Various plots. A. Plots of standard error of correlation (y-axis) versus the geometric mean heritability of the pair of traits (x-axis) for discovery (left) and replication (right) datasets. **B.** Plots of standard error of correlation (y-axis) versus the mean sample size of the pair of traits (x-axis) for discovery (left) and replication (right) datasets. **C.** Plots of difference between correlations (r_g_-r_p_, y-axis) versus the geometric mean heritability of the pair of traits (x-axis) for discovery (left) and replication (right) datasets. **D.** Plots of difference between correlations (r_g_-r_p_, y-axis) versus the mean sample size of the pair of traits (x-axis) for discovery (left) and replication (right) datasets.

**Supplementary Figure 3.**
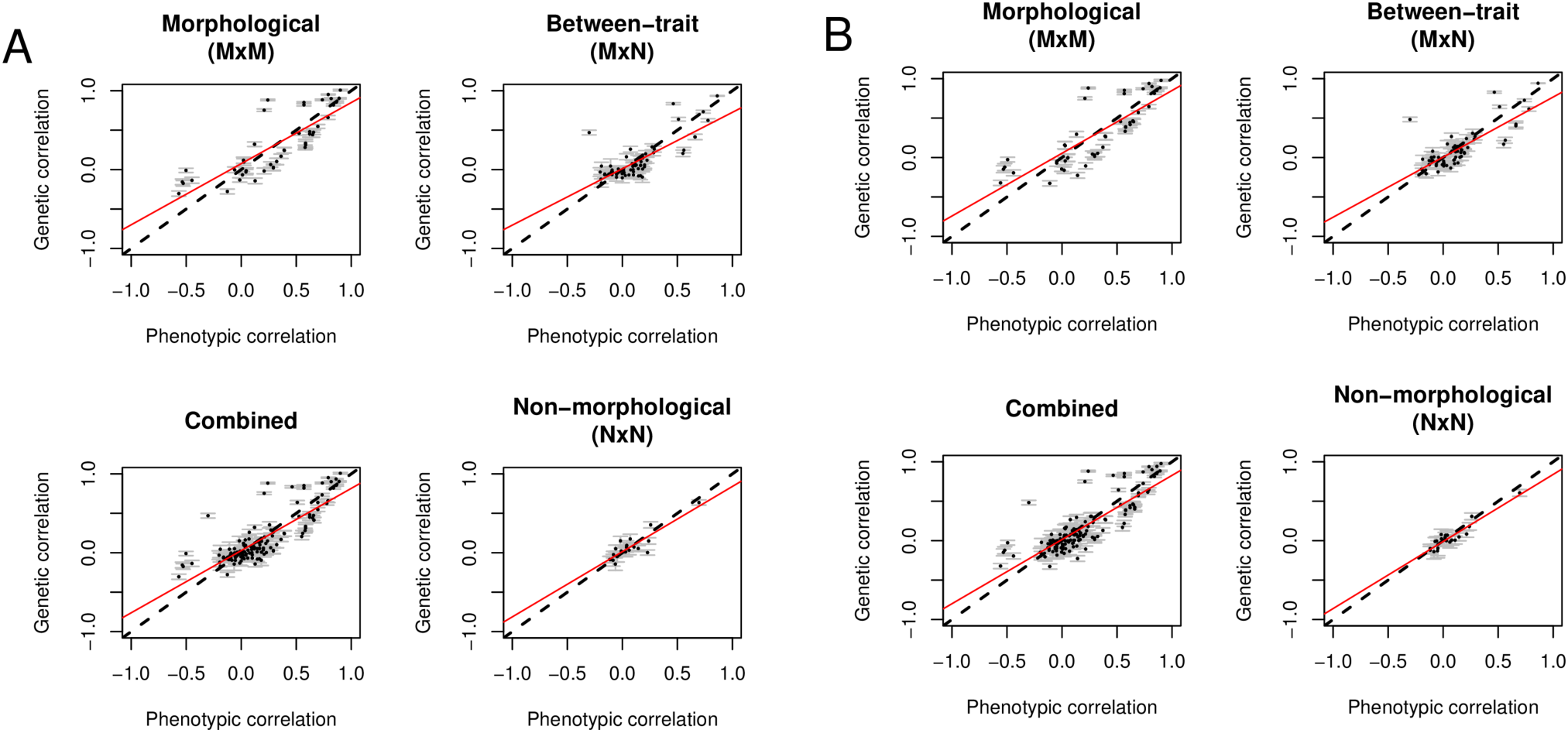
Comparison plots of between-dataset r_g_ and r_p_ without adjusting for covariates. In the main paper, the genetic correlations were calculated while adjusting for all listed covariates (Supplementary Table 2), in addition to the 15 genetically derived principal components. The phenotypic correlations did not use the 15 PCs, in order to simulate a scenario where no genetic material is available. The analysis was also run where the phenotypic correlations were calculated without covariates altogether. This figure shows the between-dataset comparison with r_g_, similar to Figure 3 presented in the main paper. Plots of genetic correlation versus environmental correlation for the between-dataset comparison. 108,035 British European individuals were evenly distributed into discovery and replication datasets. Genetic and phenotypic correlations were calculated within group for 17 traits. Environmental correlations were calculated using the formula 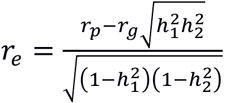, as derived in the supplementary text. **A.** Genetic correlations from discovery dataset, environmental correlations from replication dataset. **B.** Genetic correlations from replication dataset, environmental correlations from discovery dataset.

**Supplementary Table 5.** Summary statistics of linear regression (comparison of r_p_ and r_g_ without covariates) The analysis presented here was run where the phenotypic correlations were calculated without covariates. This figure shows the summary statistics of the linear regression in the between-dataset comparison with r_g_, similar to Tables 2 and 3 presented in the main paper.

|  | Discovery genetic<br>Replication environmental |  |  |  | Discovery environmental<br>Replication genetic |  |  |  |
| --- | --- | --- | --- | --- | --- | --- | --- | --- |
| | $r$ | Slope | Intercept | Avg $D_{abs}\ddagger$ | $r$ | Slope | Intercept | Avg $D_{abs}\ddagger$ |
| Morphological | 0.81*(0.06) | 0.82 | 0.04 | 0.20(0.04) | 0.83*(0.06) | 0.8† | 0.06 | 0.19(0.04) |
| Non-Morphological | 0.9*(0.07) | 0.82 | 0 | 0.07(0.07) | 0.92*(0.06) | 0.82† | 0.01 | 0.06(0.06) |
| Between-trait | 0.75*(0.08) | 0.71† | 0.02 | 0.12(0.05) | 0.81*(0.07) | 0.78† | -0.01 | 0.11(0.05) |
| Combined | 0.82*(0.03) | 0.81† | 0.02 | 0.14(0.05) | 0.85*(0.03) | 0.82† | 0.01 | 0.13(0.05) |

|  | Discovery |  |  |  | Replication |  |  |  |
| --- | --- | --- | --- | --- | --- | --- | --- | --- |
| | $r$ | Slope | Intercept | Avg $D_{abs}\ddagger$ | $r$ | Slope | Intercept | Avg $D_{abs}\ddagger$ |
| Morphological | 0.81*(0.06) | 0.82 | 0.04 | 0.20(0.04) | 0.83*(0.06) | 0.8† | 0.06 | 0.19(0.04) |
| Non-Morphological | 0.9*(0.07) | 0.82 | 0 | 0.07(0.07) | 0.92*(0.06) | 0.83† | 0.01 | 0.05(0.06) |
| Between-trait | 0.75*(0.08) | 0.7† | 0.02 | 0.12(0.05) | 0.82*(0.07) | 0.78† | -0.01 | 0.11(0.05) |
| Combined | 0.82*(0.03) | 0.8† | 0.02 | 0.14(0.05) | 0.85*(0.03) | 0.83† | 0.01 | 0.13(0.05) |

\*=significant at $p<0.0083$ (Bonferroni multiple testing correction)
†=significant difference from unity line ( $p<0.00625$ )
‡=average of absolute disparity

**Supplementary Table 6.**
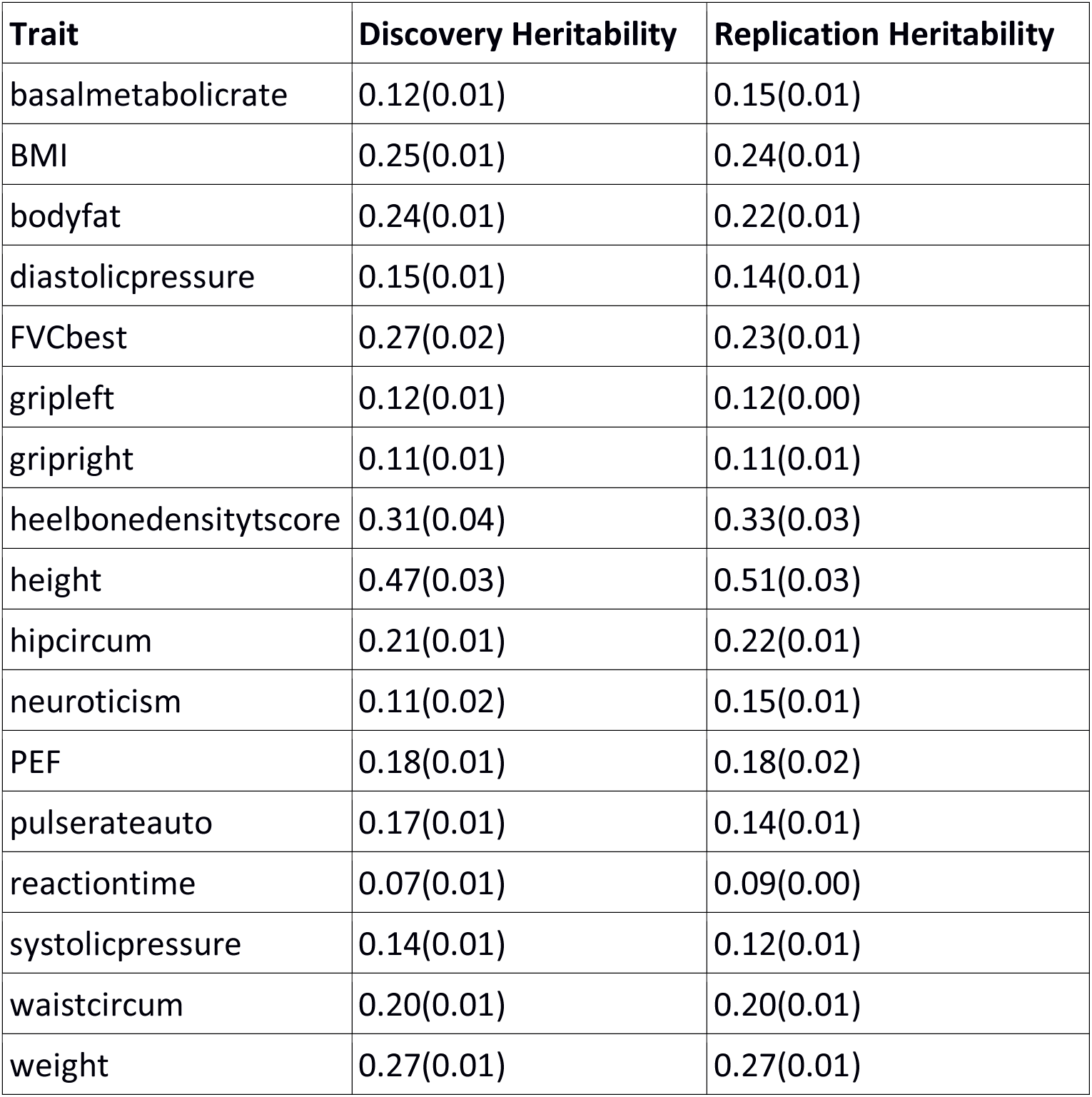
Heritability estimates with standard errors. 108,035 British European individuals were evenly distributed into discovery and replication datasets. A GWAS study was performed and LD-scores calculated. These were used to perform an LD-score regression using the software LDSC. The estimated heritability and standard error for the estimates are shown below.

**Supplementary Table 7-14- Genetic and phenotypic correlation matrices** The genetic and phenotypic correlation matrices used in this study, as well as their corresponding standard errors, are provided in the supplementary material spreadsheet.

